# Livestock grazing is associated with seasonal reduction in pollinator biodiversity and functional dispersion but cheatgrass invasion is not: variation in bee assemblages in a multi-use shortgrass prairie

**DOI:** 10.1101/2020.07.29.226621

**Authors:** Khum Bahadur Thapa-Magar, Thomas Seth Davis, Boris C. Kondratieff

## Abstract

Livestock grazing and non-native plant species affect rangeland habitats globally. These factors may have important effects on ecosystem services including pollination, yet, interactions between pollinators, grazing, and invasive plants are poorly understood. To address this, we tested the hypothesis that cattle grazing and site colonization by cheatgrass (*Bromus tectorum*) impact bee foraging and nesting habitats, and the biodiversity of wild bee communities, in a shortgrass prairie system. Bee nesting habitats (litter and wood cover) were marginally improved in non-grazed sites, though foraging habitat (floral cover and richness) did not differ among grazed, non-grazed, or cheatgrass colonized sites. However, floral cover was a good predictor of bee abundance and functional dispersion. Mean bee abundance, richness, diversity and functional diversity were significantly lower in cattle-grazed habitats than in cheatgrass-colonized or non-grazed habitats. Differences in bee diversity among habitats were pronounced early in the growing season (May) but by late-season (August) these differences eroded. Fourth-corner analysis revealed that sites with high floral cover tended to support large, social, polylectic bees; sites with high grass cover tended to support oligolectic solitary bees. Both cattle-grazed and cheatgrass-colonized sites were associated with lower abundances of above-ground nesting bees but higher abundance of below-ground nesters. We conclude that cheatgrass-invaded sites are not associated with reduced bee biodiversity or abundance, but cattle grazing was negatively associated with bee abundances and altered species composition. Although floral cover is an important predictor of bee assemblages, this was not impacted by grazing and our suggests that cattle likely impact bee communities through effects other than those mediated by forbs, including soil disturbance or nest destruction. Efforts aimed at pollinator conservation in prairie habitats should focus on managing cattle impacts early in the growing season to benefit sensitive bee species.

## Introduction

Wild bees play key functional roles in natural landscapes including the pollination of wild plants and crops and are vital for maintaining biodiversity and ecosystem function (Kearns et al. 1998, Kremen et al. 2007). Roughly 90% of the world’s plant species are pollinated by animals, in which bees are the dominant flower visitors for pollination services (Ollerton et al. 2011). However, wild bees are declining globally, with serious implications for human food security and ecosystem function (Potts et al. 2010, Vanbergen and Pollinators Initiative 2013). Most authors now agree that wild bees are vital for pollination services in agricultural systems and can exceed the services provided by honey bees (*Apis mellifera* L.) (MacInnis and Forrest 2019, Greenleaf and Kremen 2006). Accordingly, conservation of wild bee communities is important to maintain pollination services in both agricultural areas and natural landscapes.

Habitat alteration and exotic species introduction are hypothesized to be among the major contemporary drivers directly and indirectly affecting bee communities (Rafferty 2017). In rangeland ecosystems, managed livestock grazing is a dominant process by which habitat alteration occurs (Alkemade et al. 2013). Livestock grazing can impact wild bees directly or indirectly through various mechanisms, including effects on bee nesting and foraging habitats (Moreira et al. 2019) and behaviors (Sjödin 2007). For example, soil compaction due to livestock activity can damage potential or existing ground nesting sites crucial for ground- and cavity-dwelling wild bee species (Murray et al. 2012) or livestock may consume or alter composition of forbaceous species that wild bees rely on for foraging resources (Carvell 2002, Roulston and Goodell 2011). In addition, livestock may directly kill adult bees as well as their larvae via trampling (Sugden 1985, Sjödin et al. 2007). Since ground-nesting solitary bee species comprise a substantial proportion of many wild bee communities, these effects are a serious concern for ranch managers concerned with maintenance of ecosystem services and may ultimately affect rangeland productivity. In addition, repeated pressure on plant communities from livestock grazing can also impact plant growth, architecture (Kruess and Tscharntke 2002a, b), floral traits, plant attractiveness to pollinators, plant reproductive success (Jones and Agrawal 2017, Bauer et al. 2017), and soil characteristics (Potts et al. 2005). Yet, an understanding of these collective effects on wild bee pollinators in rangelands remains nascent.

In addition to managed livestock grazing, biological invasion is another ecological process driving habitat alteration in rangeland systems and may also have consequences for wild bee communities (Kearns et al. 1998). Both invasive forbs and grasses affect wild bee communities indirectly through impacts on native plant composition and abundance. Invasive plants may outcompete native forbs for nutrients, light, space and water (Levine et al. 2003, Parkinson et al. 2013). Invasive grasses, particularly *Bromus* species including *B. tectorum* L. and *B. japonicus* Thunb. (hereafter, ‘cheatgrasses’) have extensively occupied many rangeland ecosystems in western North America (Goergen et al. 2011). Invasion of rangeland habitats by cheatgrasses may impact wild bee communities via multiple mechanisms, but these interactions have not yet been examined.

To provide new information on the interactions between pastoral land use, habitat degradation via invasive species, and wild bee communities, we ask the question “Does livestock grazing (cattle) or site occupancy by cheatgrass impact bee biodiversity relative to non-grazed, non-invaded sites?” To answer this question, our objectives were to (1) compare wild bee nesting and foraging resources in rangeland habitats utilized for cattle grazing, invaded by cheatgrass, and non-grazed, non-invaded habitats; (2) analyze the relationships between these three habitat types and seasonal variation in bee assemblages and functional dispersion, and (3) characterize associations between foraging and nesting resources and bee functional traits. Our studies provide new insights into the relationship between wild bee communities and dominant ecological processes affecting their habitats in a shortgrass prairie ecosystem, with implications for the management of rangelands and maintenance of pollination services.

## Methods and materials

### Study area and description of sites

Study sites were selected in semiarid shortgrass-steppe habitats of the Front Range of Colorado. Sites were typically predominated by blue gramma (*Bouteloua gracilis* (Willd. ex Kunth) Lag. ex Griffiths) and buffalo grass (*B. dactyloides* (Nutt.); Burke et al. 2008). The shortgrass-steppe has an evolutionary history of ungulate grazing by bison and elk that predates European settlement. Following European settlement, these rangelands have been managed primarily for cattle grazing (Cook and Redente 1993). Public land management agencies in the region typically use fenced enclosures to control cattle grazing, and we took advantage of these enclosures to select study sites that were actively managed for cattle (hereafter referred to as ‘grazed’ sites); in grazed sites mean stocking rates were 93±11 (SE) animal unit months (AUM’s). Adjacent landscapes have been designated as natural areas or open spaces and set aside for conservation and recreation. These conservation areas are protected from cattle grazing but may also become heavily colonized by cheatgrasses (Banks and Baker 2011). We exploited non-grazed conservation units as a natural experiment for exclusion of cattle grazing to establish study sites that were non-grazed and exhibited minimal colonization by cheatgrasses (hereafter, ‘non-grazed’ sites) and sites that were non-grazed but had high incidence of cheatgrass colonization (hereafter, ‘cheatgrass’ sites). We selected a total of thirty sites with each habitat type (grazed, cheatgrass, or non-grazed) represented by ten sites each, and sites were separated by a minimum distance of 1 km.

### Field cover sampling

At each study site, a central point was established from which five equidistant 50 m transects originated; transects were oriented to 0°, 72°, 144°, 216° and 288° and along each transect a pin drop was used to sample cover at one meter intervals (250 total pin drops per site). Pin drops were grouped into six possible categories including rock, bare ground, wood or litter material, native grass, cheatgrass, or forb. Forbs were further characterized to have either active floral displays (i.e., flowering) or no active floral displays (not flowering or in dry-down). Transect data were aggregated to compute the proportion of ground cover that fell into each category and floral cover was determined by analyzing the proportion of forb contacts that were actively flowering at the time of sampling. Forbs actively flowering at the time of sampling were also identified in the field to the lowest possible taxonomic level to estimate richness of floral cover. To account for seasonal variation in bee foraging habitat (floral cover and richness), floral sampling was repeated four times during the growing season of 2018 in May, June, July, and August with each sampling occurring mid-month.

### Bee collection procedures

Bees were collected from each study site using a passive trapping method (‘blue vane’ traps). Traps consisted of an ultra-violet reflective blue vane fixed to a yellow collection bucket (SpringStar, Woodinville, WA, USA). Traps were placed at the previously established central location at each site to sample bee assemblages over four separate periods (May, June, July, and August) that corresponded with the assessments of floral cover. In each trapping period, traps were hung from wooden stakes at a height of 1 m, and trap contents were collected after 48 h. Bees were collected into plastic bags, placed on dry ice, and immediately returned to the laboratory for curation.

All collected bee specimens were pinned, mounted, sorted to morphospecies and were subsequently identified to the lowest taxonomic level possible, in most cases this was to genus and species. Specimen identifications were confirmed by insect taxonomists external to the study (V. Scott and A. Carper, University of Colorado; Scott et al. 2011). Vouchers of identified bee specimens are curated at the C.P. Gillette Museum of Arthropod Diversity at Colorado State University (Fort Collins, Colorado).

### Bee functional traits

Bee qualitative and quantitative functional traits were compiled for the purposes of calculating functional dispersion, a metric that describes the relative diversity of functional traits in a species assemblage (Laliberté and Legendre 2010). We considered multiple ecological traits related to wild bee life history, behavior, and foraging ranges including diet breadth (lecty), nesting habit and nest locations, pollen carrying structures, sociality, and body size (Michener 2007).

Traits including intertegular distance (ITD, a proxy for body size) and tibial hair density were resolved using high-resolution photographic methods as follows: photographs were taken for ten replicate specimens (5 male, 5 female) per species from three orientations (head, dorsal and ventral views) for each of 49 species using Canon-EOS Rebel T7i DSLR (49 species ×3 orientations ×10 specimens per species=1470 photograph images). ITD was measured from photograph layers using the image J program (Schneider et al. 2012) to generate an average value for each species. For categorical life history traits, we used scientific literature, online databases, books and field observations for traits classification (Table S1 and S2; Michener 1999, 2007, Scott et al. 2011, Cariveau et al. 2016, Danforth et al. 2019, Hall et al. 2019). Individuals that were not positively identified to species, but able to be identified to genus, were assigned trait values from the closest congener considered to have a similar life history (Michener 2007). Flight phenology (early, middle, or late-season) was assigned based on the collection period in which abundances were maximized for a given species (Table S3).

### Data analysis

All analyses were implemented in R version 3.6.2 and, unless otherwise stated, incorporate a Type I error rate of α=0.05 for assigning statistical significance. However, modeled effects were interpreted as marginally significant at the α=0.10 level. In parametric analyses using continuous variables, response and predictor variables were standardized to meet assumptions of normality and homogeneity.

#### Computation of bee diversity indices and functional dispersion

A bee species abundance matrix was used to derive species-level abundances as well as bee species richness and α-diversity (Shannon’s H’) for each site × collection date combination. We computed functional dispersion (FDis) for bee assemblages at each site × collection date combination using the methods of Laliberté and Legendre (2010) and the metrics shown in Table S1; FDis was computed using the R add-on package ‘FD’ (Laliberté et al. 2015) and applying the Cailliez correction for non-Euclidean distances generated by inclusion of categorical traits. The metrics of bee species abundance, species richness, diversity, and FDis were used as response variables in the analyses described below.

#### Objective 1: compare bee nesting and foraging resources in rangeland habitats utilized for cattle grazing, invaded by cheatgrass, and non-grazed, non-invaded habitats

The effects of site classification (n=3; ‘cattle-grazed’, ‘cheatgrass-colonized’, or ‘non-grazed’) on foraging and nesting resources was tested using one-way ANOVA. Response variables included percent of ground area characterized as rock, wood and litter, native grass, cheatgrass, bare soil, or floral cover. Floral richness was also analyzed as a response variable, and Tukey’s HSD test was used to make all pairwise comparisons among sample means. For this analysis, mean floral cover and richness were used (i.e., averaged across each month of survey). We considered bare soil cover and woody/litter material cover to be proxies for nesting substrates, and floral cover and richness as proxies for foraging habitat. Rock, grass, and cheatgrass cover are representative of cover types that are not usable by bees but were abundant in the survey areas.

#### Objective 2: analyze the relationships between habitat types and seasonal variation in bee assemblages and functional dispersion

We examined how cattle grazing or cheatgrass colonization affect bee diversity using several statistical approaches. First, we tested the fixed effects of site classification (n=3) and collection period (n=4; May, June, July, and August) and the site classification × collection period interaction on the responses of mean bee abundance, richness, diversity, and FDis using a two-way ANOVA model.

Sampling curves were generated to estimate and compare rates of species detection across the three different site classifications and was implemented using the R add-on package ‘iNEXT’ (Hsieh et al. 2016). To quantify β-diversity and turnover in genera across collection periods and sample locations, we used nonmetric multidimensional scaling (NMDS) of Bray-Curtis dissimilarities to evaluate variability in bee community assemblages across habitats and sample month.

#### Objective 3: characterize associations between foraging and nesting resources and bee functional traits

We also examined how variation in foraging and nesting resources affected bee community metrics to determine whether efforts to manage cover would have potential impacts on bee assemblages. We used a generalized linear model with an identity link function to analyze variation in bee assemblage abundance, richness, diversity, and FDis due to variation in cover composition (rock, bare soil, wood/litter, grass, cheatgrass, and floral cover) and floral richness.

To analyze the associations between specific bee functional traits and local habitat factors we used fourth-corner analysis (Legendre et al. 1997, Brown et al. 2014) implemented in the R add-on package ‘mvabund’ (Wang et al. 2020). Generalized linear models of fourth-corner statistics were fit for bee species abundances as a function of a matrix of species traits and environmental variables (and their 2-way interaction) using a Least Absolute Shrinkage and Selection Operator’s (LASSO) penalty which restricts influences of interactions that do not add to the Bayesian Information Criteria (BIC). Analysis of model deviance was estimated using a Monte-Carlo resampling procedure (9,999 resamples) to evaluate the global significance of trait-environment relationships.

## Results

### Objective 1: compare bee nesting and foraging resources in rangeland habitats utilized for cattle grazing, invaded by cheatgrass, and non-grazed, non-invaded habitats

As expected, there was significant variation in the proportion of mean cheatgrass cover (*P*<0.05) and mean native grass cover (*P*<0.05) among the site classifications, with the highest proportion of cheatgrass cover in cheatgrass-colonized sites and the highest proportion of native grass cover in cattle-grazed sites. Cattle-grazed and non-grazed sites both had low levels of cheatgrass cover and were not significantly different from one another. Mean litter and wood cover differed marginally among site classifications (*P*<0.10) and trended towards being highest at non-grazed sites. In contrast, bare soil cover, rock cover, floral cover, and floral richness were all similar among site classifications (*P*>0.05; Table 1).

**Table 1.** A comparison of cover classes and floral richness across grazed, cheatgrass, and non-grazed rangeland sites. Values are means plus or minus one standard error of the mean, and superscript lettering denotes Tukey’s HSD test.

| Variable | Cattle-grazed | Cheatgrass-colonized | Non-grazed | *F* (2,27) | *P* |
| --- | --- | --- | --- | --- | --- |
| Grass cover (native, %) | 41.4<sup>a</sup>± 2.8 | 13.9<sup>b</sup>±1.2 | 30.7<sup>a</sup>±2.0 | 23.5 | <0.001 |
| Cheatgrass cover (%) | 5.5<sup>a</sup>±1.4 | 26.3<sup>b</sup>±3.5 | 2.1<sup>a</sup>±0.6 | 18.6 | 0.003 |
| Floral cover (%) | 8.2<sup>a</sup>±3.7 | 10.2<sup>a</sup>±3.1 | 10.39<sup>a</sup>±4.1 | 0.2 | 0.890 |
| Litter/wood cover (%) | 12.5<sup>a</sup>±2.2 | 10.7<sup>a</sup>±1.4 | 18.9<sup>a</sup>±2.2 | 2.6 | 0.090 |
| Bare ground cover (%) | 12.3<sup>a</sup>±1.9 | 5.8<sup>a</sup>±1.1 | 8.84<sup>a</sup>±1.4 | 2.4 | 0.110 |
| Rock (%) | 1.1<sup>a</sup>±0.5 | 1.9<sup>a</sup>±0.6 | 0.4<sup>a</sup>±0.2 | 1.3 | 0.291 |
| Floral richness | 16.0<sup>a</sup>±0.7 | 19.0<sup>a</sup>±0.5 | 19.0<sup>a</sup>±0.5 | 0.7 | 0.475 |

### Objective 2: analyze the relationships between habitat types and seasonal variation in bee assemblages and functional dispersion

A total of 4,368 bees representing four families (Apidae, Colletidae, Halictidae, and Megachilidae) were captured in blue vane traps. The four families were represented by 18 genera and 49 species. The European honeybee, *Apis mellifera*, represented only ~2% of the total collection, indicating that cultured bees had relatively little impact on the study. Three genera including bumble bees (*Bombus* spp.), long-horned bees (*Melissodes* spp.), and furrow bees (*Halictus* spp.) collectively comprised about 63% of the sample (Table 2). Rarefaction analysis indicated that rates of species detections were similar among the three habitat classifications (Figure S1).

**Table 2.**
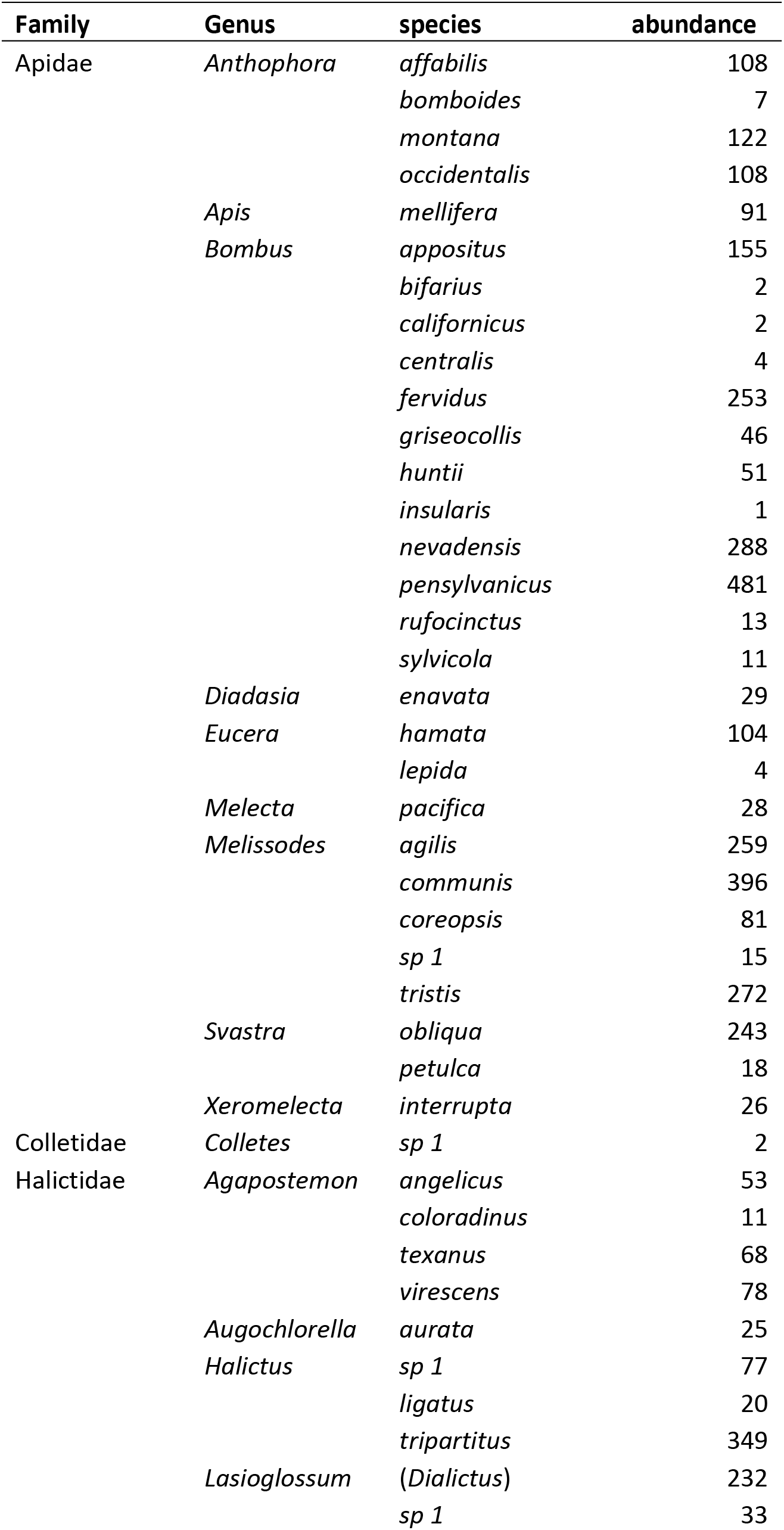
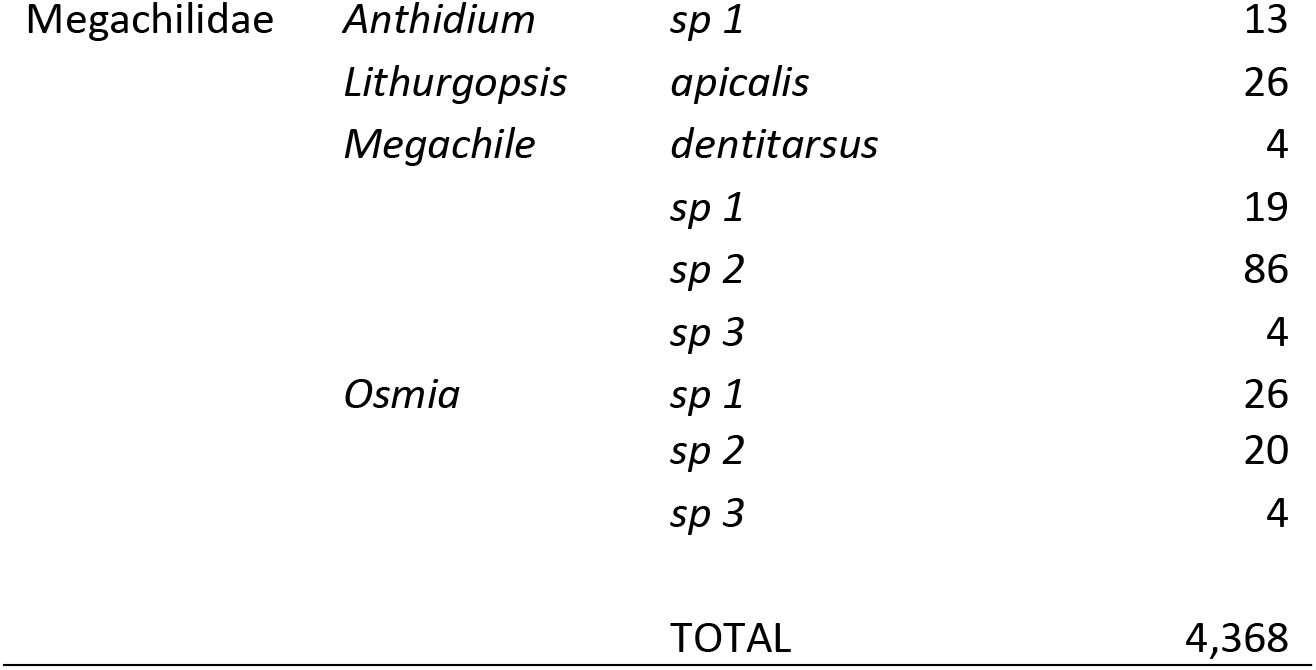
Summary of all bee taxa captured during the study and their abundances.

There were significant differences in bee community metrics due to site classification, month of collection, and their interaction. Bee abundance varied significantly due to the main effects of site classification (*F*_2, 100_=3.945, *P*=0.022) and collection period (*F*_3,100_=18.116, *P*<0.001), but there was no evidence of an interaction between these terms (*F*_6, 100_=0.846, *P*=0.537; Figure 2A). Similarly, bee species richness varied due to the main effects of site classification (*F*_2, 100_=9.293, *P*<0.001) and collection period (*F*_3,100_=23.227, *P*<0.001), but not their interaction (*F*_6,100_=1.037, *P*=0.406; Figure 2B). Bee diversity also varied significantly due to the main effects of site classification (*F*_2, 103_=10.805, *P*<0.001), collection period (*F*_3,103_=21.485, *P*<0.001), and their interaction (*F*_6,103_=2.529, *P*=0.025; Figure 2C). FDis of bee assemblages varied due to the main effect of site classification (*F*_2,100_=15.019, *P*<0.001) but did not differ across collection periods (*F*_3,100_=2.034, *P*=0.113); there was marginal evidence of an interaction between collection period and site classification (*F*_6,100_=2.048, *P*=0.066, Figure 2D).

**Figure 1.**
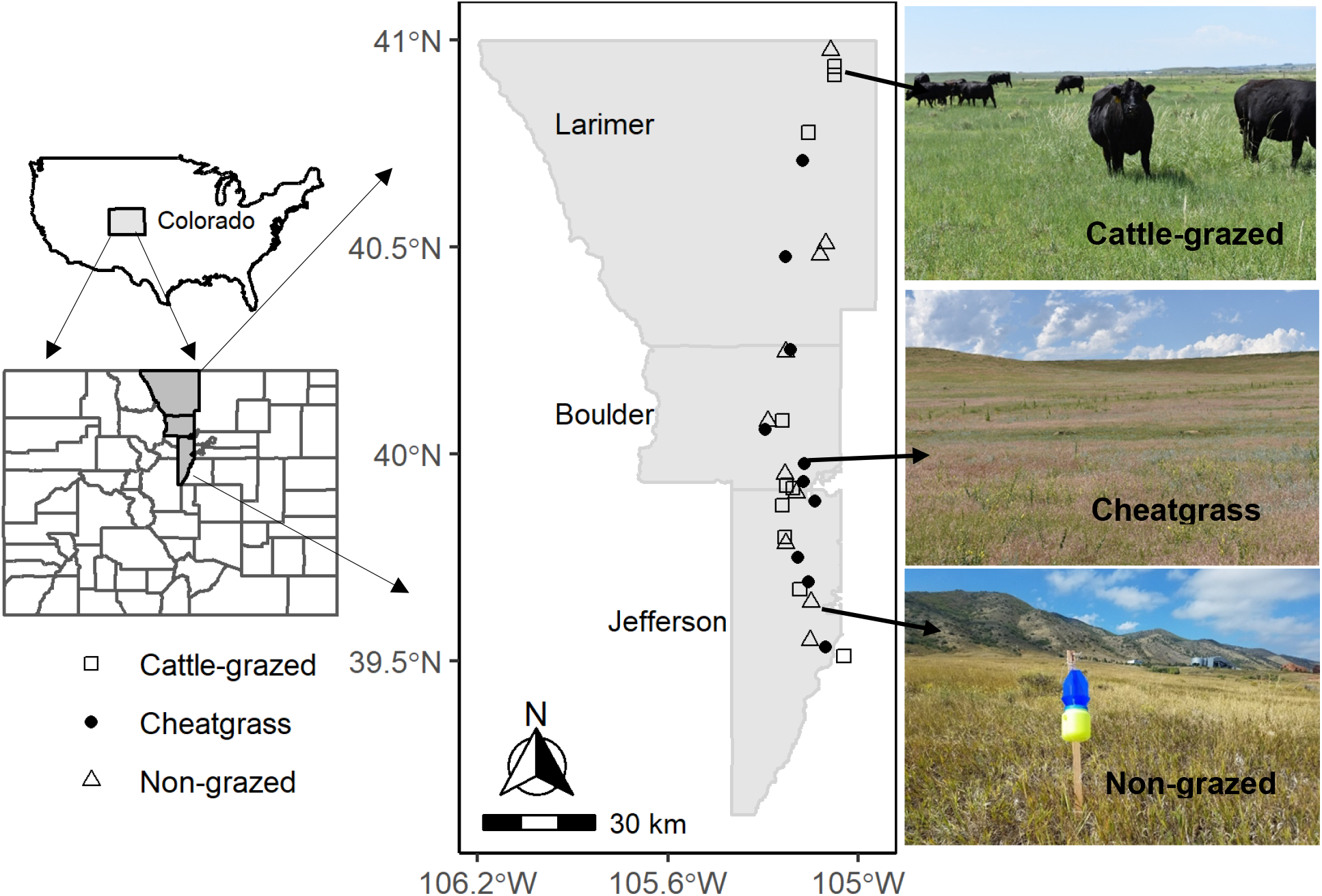
Approximate location of 30 shortgrass prairie study sites distributed across the Colorado Front Range and representative photographs of sites. Study locations were comprised of cattle-grazed sites, sites heavily colonized by cheatgrass (*Bromus* spp.), and sites that were non-grazed and with minimal cheatgrass cover.

**Figure 2.**
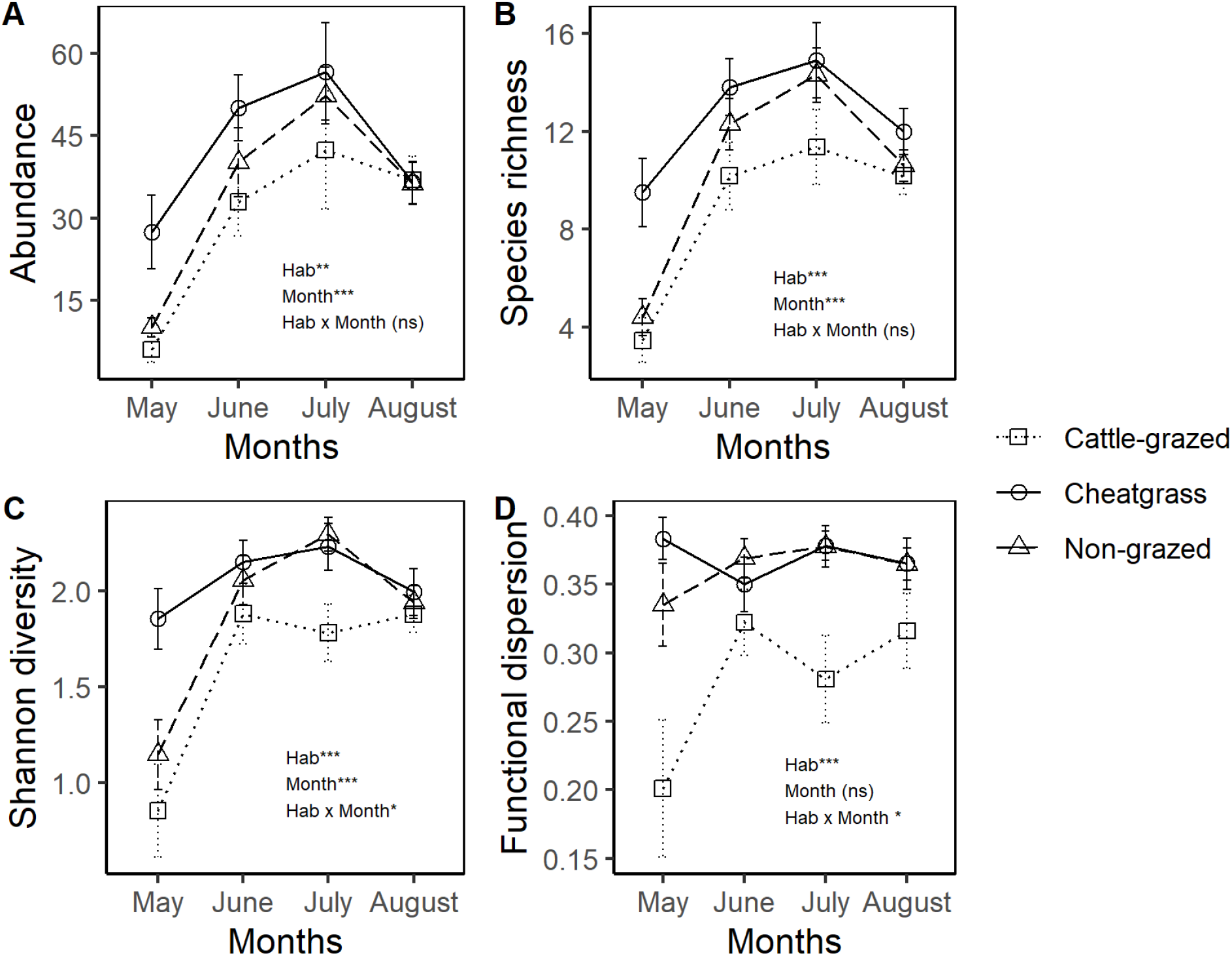
Variation in mean (A) bee abundance, (B) species richness (C) diversity, and (D) FDis represented as a habitat classification × collection period interaction.

Analysis of bee community composition with NMDS indicated distinct differences in species assemblages between cattle-grazed and non-grazed sites, but species assemblages in cheatgrass-invaded sites were similar to both grazed- and non-grazed sites (Figure 3A). Differences in species assemblages between cattle-grazed and non-grazed sites were generally reflected by a turnover in the ratio of *Bombus*: *Melissodes* species; however, abundances of multiple genera were consistent across site classification (Table S4). There were also distinct seasonal differences in the genera composition of bee assemblages with both *Bombus* and *Melissodes* becoming more abundant throughout the season and all other species generally becoming less prevalent (Figure 3B), though some genera such as *Agapostemon* were consistent in their abundances throughout the growing season (Table S4).

**Figure 3.**
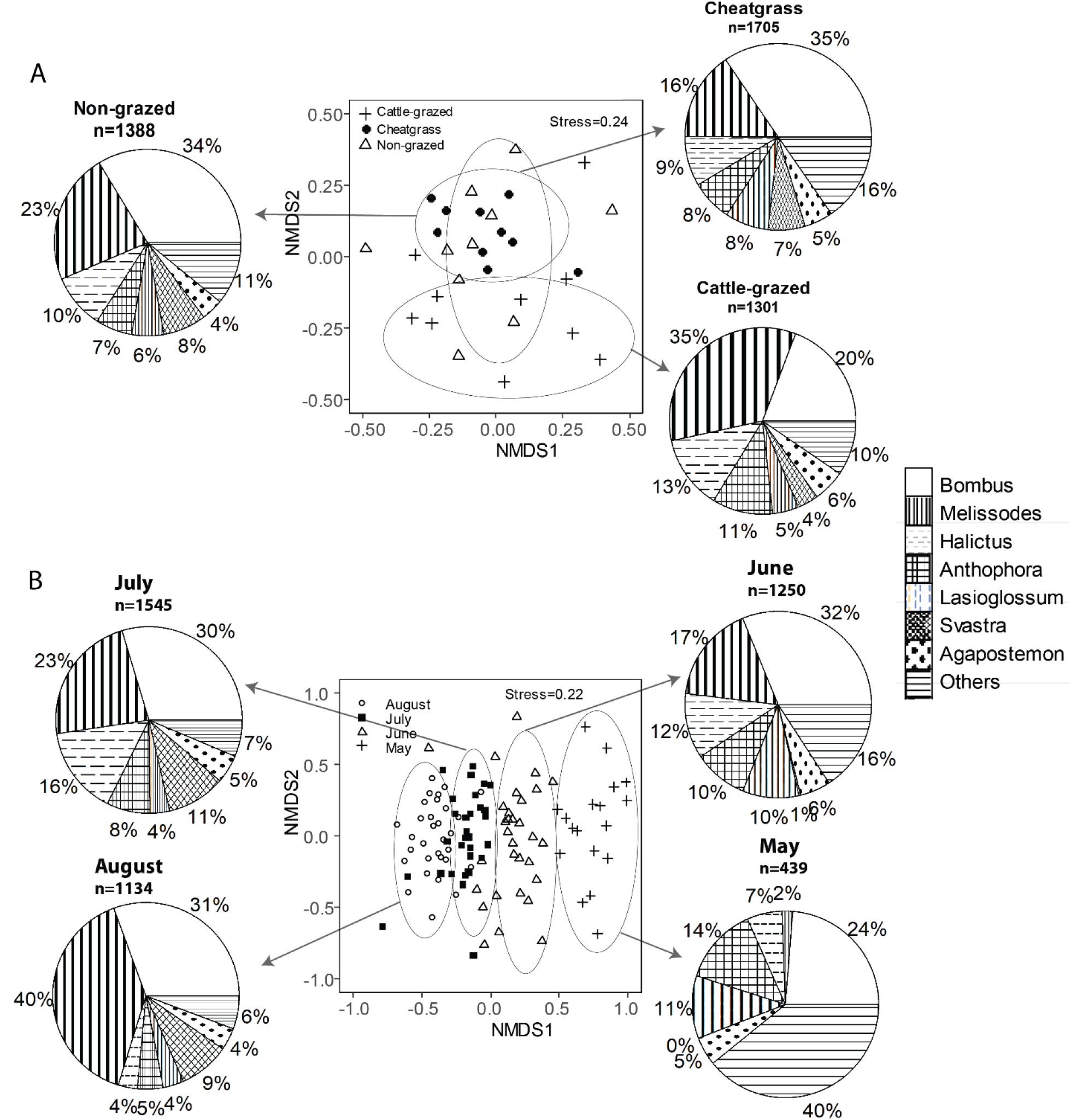
Non-metric multidimensional scaling (NMDS) of bee assemblages relative to (A) habitat classification and (B) collection period.

### Objective 3: characterize associations between foraging and nesting resources and bee functional traits

Linear model analysis testing ability of habitat components (cover) to predict variation in bee community assemblages revealed that, although elements of foraging or nesting habitat were not strongly differentiated by site classifications, some were nonetheless good predictors off bee community metrics (Table S5). Specifically, there was significant positive association between bee abundances and floral cover (β=0.549, *P*=0.037), although the species richness of bee assemblages was not associated with any cover factor or floral richness. Similarly, diversity of bee assemblages was not significantly associated with any cover factors. However, the FDis of bee communities was significantly negatively associated with increasing bare ground cover (β=−0.673, *P*=0.007), and FDis was also marginally negatively associated with increasing grass cover (β=−0.848, *P*=0.066; Figure 4).

**Figure 4.**
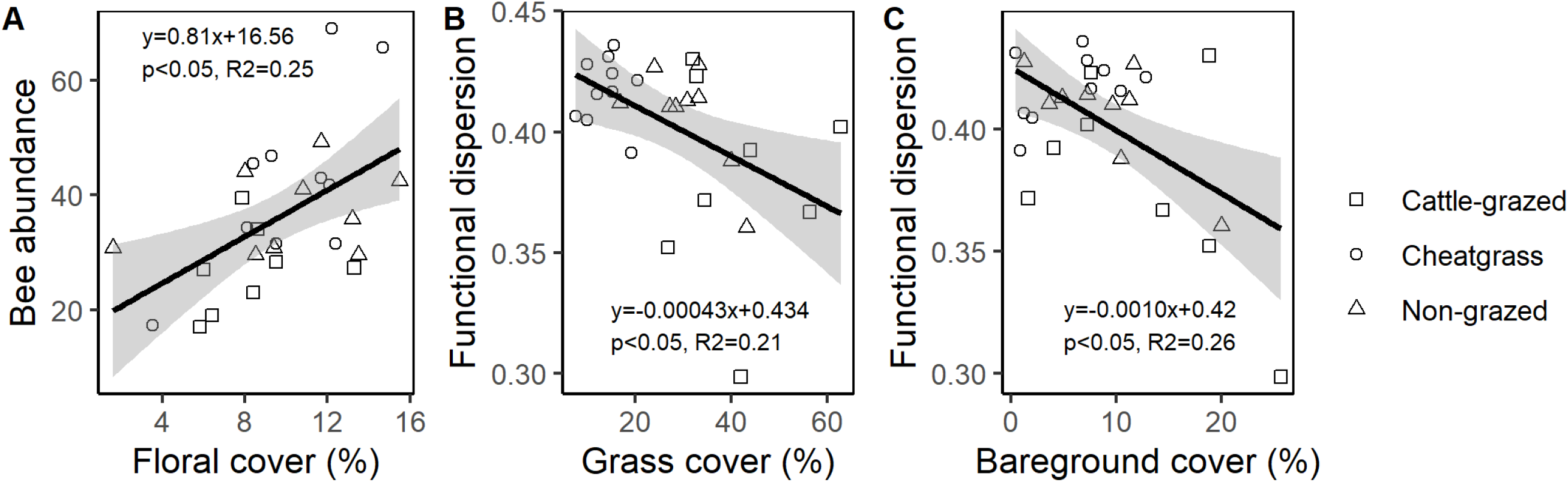
(A) Floral cover is associated with increases in bee abundances, but both (B) grass cover and (C) bare soil cover are associated with reduced functional dispersion in bee assemblages. Gray shading shows 95% confidence intervals.

Fourth-corner analysis revealed significant patterns in the correlations between habitat characteristics, bee life history traits, and bee species abundances (model deviance = 3.377, *P*<0.001). Bee body size (ITD) was positively associated with floral richness, indicating that captured bees tended to be larger as floral richness increased. Bee nest locations were correlated with habitat classification, and below-ground nesters were more abundant in cattle-grazed and cheatgrass-colonized, whereas above-ground nesters were less abundant in these areas. Diet breadth was also correlated with environmental conditions and oligolectic bees were less abundant when floral cover was high but more abundant with high grass cover, whereas the opposite was true for polylectic species; kleptoparasitic bee abundances were unrelated to cover or habitat classification. Solitary bees were less abundant in areas where floral cover and richness was high but increased in abundance in areas with high grass cover and bare soil, whereas social species were more abundant with increasing floral richness but were negatively associated with grass and bare soil cover. Variation in abundances of kleptoparasitic species and species with flexible social behaviors were not related to cover or habitat classification. Only bee species exhibiting early-season phenologies were impacted by cover, and early-season species were more abundant in areas colonized by cheatgrass. Abundances of bee species also varied due to interactions between pollen collection-related traits and environmental conditions. Bees with scopa pollen collection structures were positively associated with high grass and soil cover but negatively associated with high floral richness and rock cover, whereas bees with corbicula were positively associated with high floral richness and rock cover but negatively associated with cheatgrass and bare soil cover. Abundances of kleptoparasitic species and those with abdominal corbicula were unrelated to variation in cover or habitat classification. Variation in tibial hair densities had complex relationships with environmental conditions; bees with high tibial hair densities were more abundant in areas with high grass and soil cover, whereas bees with low tibial hair densities were more abundant in areas with high floral richness and rock cover, and bees with intermediate tibial hair densities were most abundant in areas with high floral and cheatgrass cover (Figure 5).

**Figure 5.**
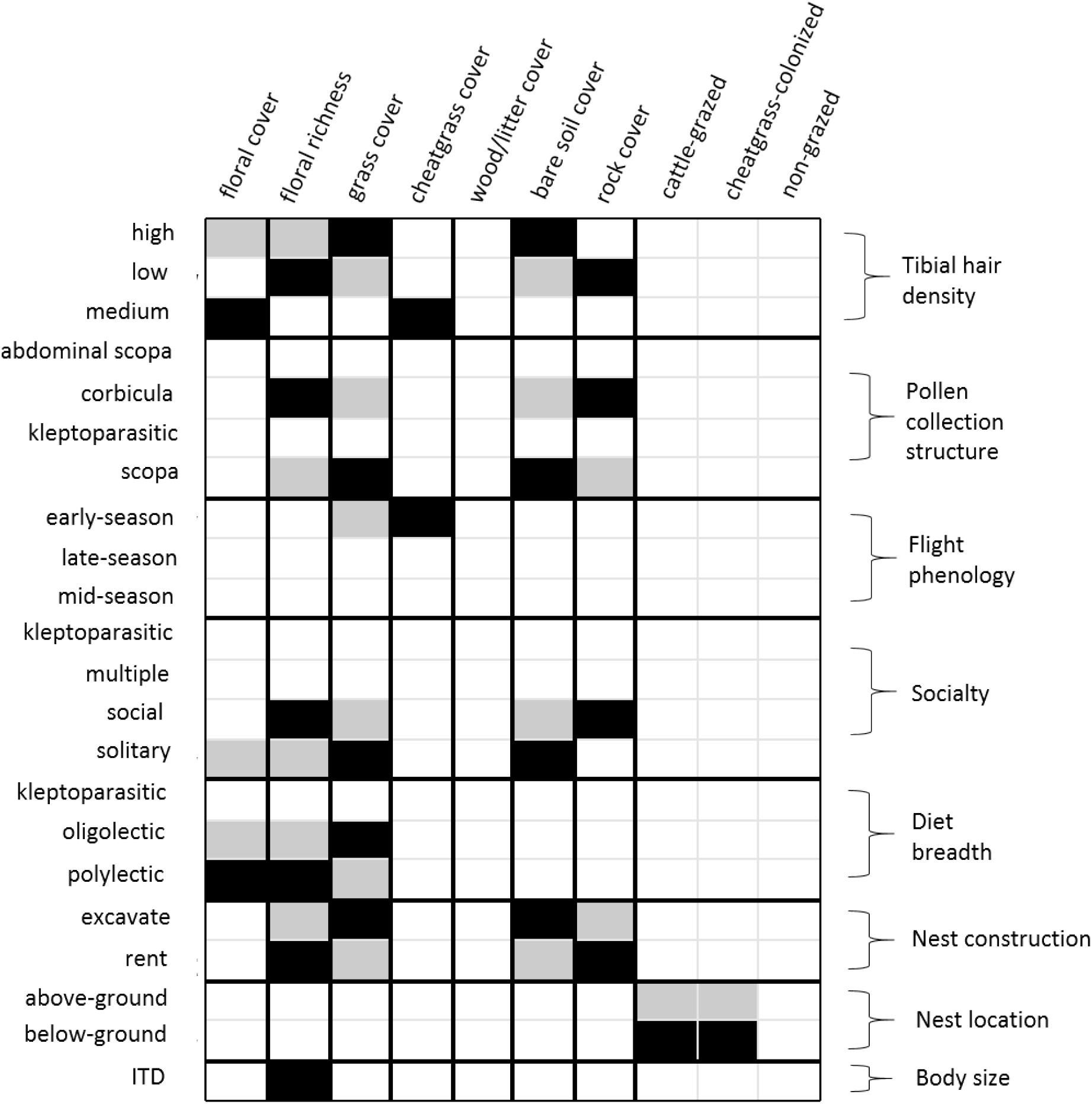
Summary of fourth-corner analysis to model bee species abundances as a function of life history trait × environment interactions. Black cells indicate positive regression coefficients, gray cells indicate negative coefficients. Blank cells indicate no relationship. Identified associations are significant at *P*<0.10.

## Discussion

Cattle-grazing and cheatgrass colonization of shortgrass prairie sites were not associated with large differences in bee foraging habitats (floral cover and species richness) but did reflect a difference in wild bee nesting habitats in terms of the proportion of native grass cover and woody material on the ground surface (Table 1). Despite the modest differences in cover composition across habitat classifications there was evidence that cattle grazing is associated with significant reductions in early- and mid-season bee diversity and FDis, but cheatgrass cover was not (Figure 2). There were distinct differences in community composition between cattle-grazed and non-grazed sites that was reflected by turnover in the ratio of *Bombus* spp: *Melissodes* spp.; however, bee assemblages in cheatgrass sites were similar to both grazed and non-grazed locations and were the most variable overall (Figure 3). Collectively, these results indicate that FDis in bee communities is more strongly predicted by broad-scale habitat classification (i.e., cattle-grazed vs. cheatgrass-colonized vs. non-grazed) than cover composition within specific sites, with potential consequences for pollination services in rangelands.

Landscapes in the study region share a long evolutionary history with bison, elk, and other wild grazing and browsing species (Milchunas et al. 1988) and forbs may therefore be well-adapted to tolerate grazing, which could partially explain why no differences in floral cover were observed across site classifications. Nonetheless, floral cover predicted bee abundances with more bees captured from sites with abundant flowering forbs (Figure 4). In other recent studies, locations with high floral density have been associated with fewer bee captures in passive traps (e.g., Rhoades et al. 2017) due to reduced attractiveness of traps when abundant floral resources are available.

Analysis of bee functional traits relative to floral cover and richness revealed that the preponderance of bees at sites with high floral cover were those with life history traits that included sociality, polylecty, and large body size. In our collections, this combination of traits is mostly represented by bumblebees (*Bombus* spp.). Accordingly, management efforts aimed at increasing or restoring local floral densities may be more likely to benefit *Bombus* spp. than other taxa. Interestingly, site colonization by cheatgrass and cattle-grazed had similar relationships with bee functional traits and were positively associated with higher abundances of bees with below-ground nesting habits (Figure 5). In some landscapes cattle may trample sensitive arthropod species resulting in reduced abundances (Bonte and Maes 2008), but this does not appear to be the case for below-ground nesting bees in our system. Although bee abundances did not differ between grazed and non-grazed sites, cattle grazing was associated with significant reductions in bee FDis indicating that cattle presence may result in a loss of bee functional diversity. The mechanisms underlying this pattern merit further study, as pollination services are generally improved with increasing bee functional diversity (Martins et al. 2015). Since floral abundance and richness were not negatively impacted at grazed sites, we hypothesize that impacts of cattle on bee assemblage functional diversity are mediated via nesting habitats, rather than through indirect consumption-mediated effects on foraging habitat. In other systems cattle-grazing has been documented to have positive effects on bee abundances even at very high grazing intensities (Vulliamy et al. 2006), so it may be difficult to generalize cattle-driven effects on bee communities.

To our knowledge, this is among the first studies to evaluate the effects of a non-native grass on pollinator assemblages. Our findings suggest that sites with a high proportion of cheatgrass were not associated with significant reductions in bee abundance, diversity, of FDis; instead, cheatgrass-dominated sites tended to have higher bee abundance and diversity early in the growing season (Figure 2). This contrasts with other recent studies. For instance, Bhandari et al. (2018) determined that pollinator abundances in semi-arid pastures were reduced under high densities of non-native forage species. Several non-mutually exclusive hypotheses could potentially explain this pattern. First, it is possible that at cheatgrass-colonized sites vane traps were more visually apparent due to the relatively homogenous structure of the vegetation and thus more attractive to foraging bees. Alternatively, sites that are occupied by cheatgrass may simply be on highly productive or suitable soils; in other words, highly productive sites may be generally superior for invasive grasses, forbs, and pollinators alike. However, this seems unlikely as floral cover did not differ between site classifications (Table 1). In future studies it will be important to determine whether the effects of cheatgrass colonization are consistently associated with high early-season bee abundance and diversity and, if so, whether these effects are an artifact of sampling strategy or due to some ecological effect such as improved nesting habitat. Accordingly, our findings do not currently suggest a need to mitigate cheatgrass occurrence for pollinator conservation efforts.

Seasonal variation in wild bee assemblage richness and functional diversity were considerable, and our sample underscores the importance of making collections across the growing season to generate reliable estimates of bee richness and diversity. There was evident turnover in taxa with certain species of *Eucera*, *Melecta*, and *Osmia* prevalent early in the growing season, but by June and July *Bombus*, *Halictus*, *Lasioglossum*, and *Melissodes* were predominant in study sites (Table S3). Altogether, bee taxa richness and diversity were lowest in the early growing season, which is consistent with other reports (Rhoades et al. 2018) and was mostly due to the relative inactivity of many social and semi-social species in the spring. Our collection had a lower rate of species detection than other regional studies focusing on bees in Colorado grasslands. For example, Kearns and Oliveras (2009) detected 108 species in grasslands of Boulder County, Colorado and an earlier study by Cockerell (1907) detected 116 species. In addition, these earlier studies found that floral resources were generally positively associated with intermediate levels of cattle grazing, which was not supported in the present study. In both earlier studies, collections were continued for several years (up to 5) and using hand netting methods--which is often associated with a higher rate of species detection than passive sampling methods (Rhoades et al. 2017), though rates of species detection in netting-based collections are presumably impacted by observer bias and skill (Westphal et al. 2008). However, bee abundances in the present study were similar to those found in both earlier works. The largest effects on bee diversity and FDis occurred early in the growing season (Figure 2), potentially indicating that species active primarily in spring have behavioral or life history traits that predispose them to site disturbance by livestock.

Collectively, our results have several implications for managers concerned with maintaining site occupancy by wild bee assemblages in rangelands where livestock production is a common land use. First, our results do not suggest that floral resources are enhanced in sites managed for cattle grazing as some earlier studies do. Neither did we find any evidence that grazed sites exhibited any reduction in floral resources, likely indicating that grazing practices in the region do not strongly impact bee foraging habitats. However, floral resource availability was an important predictor of bee abundances. Second, bee assemblage composition did vary between grazed and non-grazed sites, and this was reflected by shifts in the ratios of *Bombus* spp: *Melissodes* spp. Further experimental work could help to elucidate whether this turnover in bee taxa is associated with variation in pollination services. Thirdly, both cattle grazing and cheatgrass were associated with reduced site occupancy by above-ground nesting bees but increased site occupancy by below-ground nesting bees. Fourth, cattle grazing was clearly associated with reduced FDis in early-season bee assemblages, and these effects may be mediated by cattle-driven impacts on nesting habitats rather than floral cover. Lastly, our study does not indicate that cheatgrass colonization of sites is likely to negatively impact bee abundance or diversity and may provide good nesting habitat. The mechanisms underlying this relationship are beyond the scope of the current study, but could have consequences for bee conservation, especially under widespread policies aimed at restoring cheatgrass-invaded habitats. For example, cheatgrass-dominated rangeland and forest sites are often treated with chemical (Baker et al. 2009), cultural (Cox and Anderson 2004), and physical (Young and Clements 2000) control methods with the general objective of reducing cheatgrass cover. Given that our study found an increased abundance of wild bees in cheatgrass sites, it will be important to determine whether cheatgrass control methods have deleterious, beneficial, or null impacts on bee assemblages to make appropriate management decisions about bee conservation.

## Acknowledgements

Funding was provided by McIntire-Stennis appropriations (USDA NIFA COL 00767). Samuel Murray, Fiona Horne and Haley Obermueller provided field and lab assistance. The staff of the C.P. Gillette Museum (Colorado State University) provided photography and storage/working space. We are also grateful to Virginia Scott and Dr. Adrian Carper (University of Colorado) for assistance with bee identification.

## Author contributions

KBT and TSD conceived and designed the studies. KBT collected the data, conducted data analyses and wrote the first version of the paper. BK and KBT identified bee specimens. All authors contributed to editing and approved subsequent drafts.

## Data accessibility

Data for all analyses, including transect data, species abundance matrices, and functional trait matrices will be uploaded to the Dryad Digital Repository upon acceptance of the manuscript for publication.

## References

Alkemade, R., Reid, R. S., van den Berg, M., de Leeuw, J., & Jeuken, M. (2013). Assessing the impacts of livestock production on biodiversity in rangeland ecosystems. Proceedings of the National Academy of Sciences of the USA 110:20900–20905.

Baker, W.L., Garner, J., & Lyon, P. (2009). Effects of imazapic on cheatgrass and native plants in Wyoming big sagebrush restoration for Gunnison sage-grouse. Natural Areas Journal 29:204–209.

Banks, R. E., & Baker, W. L. (2011). Scale and pattern of cheatgrass (*Bromus tectorum*) invasion in Rocky Mountain National Park. Natural Areas Journal 31:377–390.

Bauer, A. A., Clayton, M. K., & Brunet, J. (2017). Floral traits influencing plant attractiveness to three bee species: Consequences for plant reproductive success. American Journal of Botany 104:772–781.

Bhandari, K. B., West, C. P., Longing, S. D., Brown, C. P., Green, P. E., & Barkowsky, E. (2018). Pollinator abundance in semiarid pastures as affected by forage species. Crop Science 58:2665–2671.

Bonte, D., & Maes, D. (2008). Trampling affects the distribution of specialised coastal dune arthropods. Basic and Applied Ecology 9:726–734.

Brown, A. M., Warton, D. I., Andrew, N. R., Binns, M., Cassis, G., & Gibb, H. (2014). The fourth-corner solution - using predictive models to understand how species traits interact with the environment. Methods in Ecology and Evolution 5:344–352.

Burke, I. C., Lauenroth, W. K., Antolin, M. F., Derner, J. D., Milchunas, D. G., Morgan, J. A., et al. (2008). The future of the shortgrass steppe. In: Lauenroth, W. K. & Burke, I. C., eds. Ecology of the Shortgrass Steppe: A Long-Term Perspective. Oxford University Press, New York. P 484–510.

Cariveau, D. P., Nayak, G. K., Bartomeus, I., Zientek, J., Ascher, J. S., Gibbs, J., & Winfree, R. (2016). The allometry of bee proboscis length and its uses in ecology. PLOS ONE 11:e0151482. doi: 10.1371/journal.pone.0151482

Carvell, C. (2002). Habitat use and conservation of bumblebees (*Bombus* spp.) under different grassland management regimes. Biological Conservation 103:33–49.

Cockerell, T. D. A. (1907) The bees of Boulder County, Colorado. Univ Colo Stud IV (4), Boulder. *Ca*. Paper 373.

Cook, W., & Redente, E. (1993). Development of the ranching industry in Colorado. Rangelands 15:204–207.

Cox, R.D., & Anderson, V.J. (2004). Increasing native diversity of cheatgrass-dominated rangeland through assisted succession. Rangeland Ecology and Management 57:203–210.

Danforth, B. N., Minckley, R. L., Neff, J. L., & Fawcett, F. (2019). The Solitary Bees: Biology, Evolution, Conservation. Princeton University Press, New Jersey. 488 p.

Goergen, E. M., Leger, E. A., & Espeland, E. K. (2011). Native perennial grasses show evolutionary response to *Bromus tectorum* (cheatgrass) invasion. PLOS ONE 6: e18145. doi: 10.1371/journal.pone.0018145.

Greenleaf, S. S., & Kremen, C. (2006). Wild bees enhance honey bees’ pollination of hybrid sunflower. Proceedings of the National Academy of Sciences of the USA 103:3890–3895.

Hall, M. A., Nimmo, D. G., Cunningham, S. A., Walker, K., & Bennett, A. F. (2019). The response of wild bees to tree cover and rural land use is mediated by species’ traits. Biological Conservation, 231, 1–12.

Hsieh, T.C., Ma, K.H., & Chao, A. (2016). iNEXT: an R package for rarefaction and extrapolation of species diversity (H ill numbers). Methods in Ecology and Evolution 7:1451–1456.

Jones, P. L., & Agrawal, A. A. (2017). Learning in insect pollinators and herbivores. Annual Review of Entomology 62:53–71.

Kearns, C. A., Inouye, D. W., & Waser, N. M. (1998). Endangered mutualisms: the conservation of plant-pollinator interactions. Annual Review of Ecology and Systematics 29:83–112.

Kearns, C. A., & Oliveras, D. M. (2009). Environmental factors affecting bee diversity in urban and remote grassland plots in Boulder, Colorado. Journal of Insect Conservation 13:655–665.

Kremen, C., Williams, N. M., Aizen, M. A., Gemmill-Herren, B., LeBuhn, G., Minckley, R., et al. (2007). Pollination and other ecosystem services produced by mobile organisms: a conceptual framework for the effects of land-use change. Ecology Letters 10:299–314.

Kruess, A., & Tscharntke, T. (2002a). Contrasting responses of plant and insect diversity to variation in grazing intensity. Biological Conservation 106:293–302.

Kruess, A., & Tscharntke, T. (2002b). Grazing intensity and the diversity of grasshoppers, butterflies, and trap-nesting bees and wasps. Conservation Biology 16:1570–1580.

Laliberté, E., & Legendre, P. (2010). A distance-based framework for measuring functional diversity from multiple traits. Ecology 91:299–305.

Laliberté, E., Legendre, P. & Shipley, B. (2015). Package ‘FD’: Measuring functional diversity (FD) from multiple traits, and other tools for functional ecology. V.1.0-12.

Legendre, P., Galzin, R., & Harmelin-Vivien, M. L. (1997). Relating behavior to habitat: solutions to the fourth-corner problem. Ecology 78:547.

Levine, J. M., Vilà, M., Antonio, C. M. D., Dukes, J. S., Grigulis, K., & Lavorel, S. (2003). Mechanisms underlying the impacts of exotic plant invasions. Proceedings of the Royal Society of London. Series B: Biological Sciences 270:775–781.

MacInnis, G., & Forrest, J. R. K. (2019). Pollination by wild bees yields larger strawberries than pollination by honey bees. Journal of Applied Ecology 56:824–832.

Martins K. T., Gonzalez, A., & Lechowicz, M. J. (2015). Pollination services are mediated by bee functional diversity and landscape context. Agriculture, Ecosystems & Environment 200:12–20.

Michener, C. D. (1999). The corbiculae of bees. Apidologie 30:67–74.

Michener, C. D. (2007). The bees of the world, 2nd edition. The Johns Hopkins University Press, Baltimore. 953 p.

Milchunas, D. G., Sala, O. E., & Lauenroth, W. K. (1988). A generalized model of the effects of grazing by large herbivores on grassland community structure. American Naturalist 13:87–106.

Moreira, X., Castagneyrol, B., Abdala-Roberts, L., & Traveset, A. (2019). A meta-analysis of herbivore effects on plant attractiveness to pollinators. Ecology 100:e02707.

Murray, T. E., Fitzpatrick, U., Byrne, A., Fealy, R. Brown, M. J., & Paxton, R. J. (2012). Local-scale factors structure wild bee communities in protected areas. Journal of Applied Ecology 49:998–1008.

Ollerton, J., Winfree, R., & Tarrant, S. (2011). How many flowering plants are pollinated by animals? Oikos 120:321–326.

Parkinson, H., Zabinski, C., & Shaw, N. (2013). Impact of native grasses and cheatgrass (*Bromus tectorum*) on Great Basin forb seedling growth. Rangeland Ecology & Management 66:174–180.

Potts, S. G., Vulliamy, B., Roberts, S., O’Toole, C., Dafni, A., Ne’eman, G., & Willmer, P. (2005). Role of nesting resources in organising diverse bee communities in a Mediterranean landscape. Ecological Entomology 30:78–85.

Potts, S. G., Biesmeijer, J. C., Kremen, C., Neumann, P., Schweiger, O., & Kunin, W. E. (2010). Global pollinator declines: trends, impacts and drivers. Trends in Ecology & Evolution 25:345–353.

Rafferty, N. E. (2017). Effects of global change on insect pollinators: multiple drivers lead to novel communities. Current Opinion in Insect Science 23:22–27.

Rhoades, P., Griswold, T., Waits, L., Bosque-Perez, N. A., Kennedy, C. M., & Eigenbrode, S. D. (2017). Sampling technique affects detection of habitat factors influencing wild bee communities. Journal of Insect Conservation 21:703–714.

Rhoades, P. R., Davis, T. S., Tinkham, W. T., & Hoffman, C. M. (2018). Effects of seasonality, forest structure, and understory plant richness on bee community assemblage in a southern Rocky Mountain mixed conifer forest. Annals of the Entomological Society of America 111:278–284.

Roulston, T. H., & Goodell, K. (2011). The role of resources and risks in regulating wild bee populations. Annual Review of Entomology 56:293–312.

Scott, V. L., Ascher, J. S., Griswold, T., & Nufio, C. R. (2011). The bees of Colorado (Hymenoptera: Apoidea: Anthophila). Natural History Inventory of Colorado, University of Colorado Natural History Museum, Boulder. 101 p.

Schneider, C. A., Rasband, W. S., & Eliceiri, K. W. (2012). NIH Image to ImageJ: 25 years of image analysis. Nature Methods 9:671–675.

Sjödin, N. E. (2007). Pollinator behavioural responses to grazing intensity. Biodiversity and Conservation 16:2103–2121.

Sjödin, N. E., Bengtsson, J., & Ekbom, B. (2007). The influence of grazing intensity and landscape composition on the diversity and abundance of flower-visiting insects. Journal of Applied Ecology 45:763–772.

Sugden, E.A. (1985). Pollinators of *Astragalus monoensis* Barneby (Fabaceae): new host records: potential impact of sheep grazing. The Great Basin Naturalist 45:299–312.

Vanbergen, A. J., & The Insect Pollinators Initiative. (2013). Threats to an ecosystem service: pressures on pollinators. Frontiers in Ecology and the Environment 11:251–259.

Vulliamy, B. G., Potts, S. G., & Willmer, P. (2006). The effects of cattle grazing on plant-pollinator communities in a fragmented Mediterranean landscape. Oikos 114:529–543.

Wang, Y., Naumann, U., Eddelbuettel, D., Wilshire, J., Warton, D., Byrnes, J., et al. (2020). Package ‘mvabund’: Statistical methods for analyzing multivariate abundance data. V. 4.1.3.

Westphal, C., Bommarco, R., Carré, G., Lamborn, E., Morison, N., Petanidou, T., et al. (2008). Measuring bee diversity in different European habitats and biogeographical regions. Ecological Monographs 78:653–671.

Young, J.A., & Clements, C.D. (2000). Cheatgrass control and seeding. Rangelands 22:3–7.

